# Application of the immunoregulatory receptor LILRB1 as a novel crystallisation chaperone for class I peptide-MHC complexes

**DOI:** 10.1101/213207

**Authors:** Fiyaz Mohammed, Daniel H. Stones, Benjamin E. Willcox

## Abstract

X-ray crystallographic studies of class I peptide-MHC molecules (pMHC) continue to provide important insights into immune recognition, however their success depends on generation of diffraction-quality crystals, which remains a significant challenge. While protein engineering techniques such as surface-entropy reduction and lysine methylation have proven utility in facilitating and/or improving protein crystallisation, they risk affecting the conformation and biochemistry of the class I MHC antigen binding groove. An attractive alternative is the use of noncovalent crystallisation chaperones, however these have not been developed for pMHC. Here we describe a method for promoting class I pMHC crystallisation, by exploiting its natural ligand interaction with the immunoregulatory receptor LILRB1 as a novel crystallisation chaperone. First, focussing on a model HIV-1-derived HLA-A2-restricted peptide, we determined a 2.4Å HLA-A2/LILRB1 structure, which validated that co-crystallisation with LILRB1 does not alter conformation of the antigenic peptide. We then demonstrated that addition of LILRB1 enhanced the crystallisation of multiple pMHC complexes, and identified a generic condition for initial co-crystallisation. LILRB1 chaperone-based pMHC crystallisation enabled structure determination for class I pMHC previously intransigent to crystallisation, including both conventional and post-translationally-modified peptides, of diverse lengths. LILRB1 chaperone-mediated crystallisation should expedite molecular insights into the immunobiology of diverse immune-related diseases and immunotherapeutic strategies, particularly involving class I pMHC complexes that are challenging to crystallise. Moreover, since the LILRB1 recognition interface involves predominantly non-polymorphic regions of the MHC molecule, the approach we outline could prove applicable to a diverse range of class I pMHC.

## 1. Introduction

A molecular understanding of the class I MHC molecule has been pivotal in deciphering its central role in T cell immunity. Seminal studies in the 1980s established its remarkable ability to directly present a diverse repertoire of peptide antigens, typically 8-10 amino acids in length and derived from proteasomal degradation of intracellular proteins, for T cell recognition (1, 2). Recognition of specific MHC-presented peptides by T cell receptor (TCR) bearing CD8^+^ T lymphocytes results in cytotoxic responses and production of pro-inflammatory cytokines, key components of anti-viral and anti-tumour immunity. In addition, the engagement of pMHC complexes by receptors that belong to the immunoglobulin superfamily including killer cell immunoglobulin-like receptors and leukocyte immunoglobulin-like receptors (LILRs) is crucial for regulating diverse immune responses (3).

From the initial descriptions of class I MHC architecture (2), which highlighted a highly polymorphic groove containing electron density corresponding to bound antigen peptides, structural analyses of pMHC complexes, to date still predominantly focussed on X-ray crystallographic approaches, have led the way in our efforts to understand MHC function. While these have established fundamental molecular principles underlying peptide antigen presentation and T cell recognition (4) structural studies of pMHC molecules continue to provide major insights into the critical role of antigenic peptides in disease pathogenesis (5, 6), immunotherapeutic strategies (7, 8) and into poorly understood aspects of T cell recognition, such as post-translational modification (9–11).

Despite the advent of recombinant methods, availability of extended screens, introduction of crystallisation nanovolume robotics and dramatic technological advances in synchrotron radiation sources, the requirement to overcome the “crystallisation bottleneck” is still a significant impediment to such X-ray crystallographic analyses of pMHC (12). Consequently, reliably achieving structure determinations for predefined pMHC targets can be challenging, a fact exacerbated by the huge diversity of MHC alleles and antigenic peptides of interest. In addition to standard crystallisation techniques such as sparse matrix sampling and seeding techniques, a number of novel strategies are available to facilitate crystallisation of challenging proteins, including the surface-entropy reduction approach (13) involving substitution of lengthy side chains with alanine, S, H and Y), and chemical modification of lysine residues by reductive methylation (14). These clearly have proven utility but are not successful for every protein, and also have the potential to interfere with the delicate chemistry of the biologically critical class I MHC antigen-binding groove. An alternative is the use of non-covalent crystallisation protein chaperones (15). This approach involves co-crystallizing a target protein, such as an antibody fragment, and can promote crystallization by reducing target conformational heterogeneity and providing an additional surface for crystal contacts (16). While superficially appealing, it is unclear how this approach could best be applied to pMHC molecules.

This study describes a novel non-covalent crystallisation chaperone methodology to efficiently facilitate crystallisation of pMHC molecules, which exploits a natural ligand interaction involving LILRB1, an immunoregulatory receptor that binds a diverse range of classical and non-classical MHC molecules (17, 18). This strategy has been applied to both conventional and post-translationally modified pMHC complexes that were recalcitrant to crystallisation, facilitating both crystallisation and structure determination. This provides a new approach to catalyse molecular studies of immunobiologically important pMHC complexes.

## 2. Materials and Methods

### 2.1 Cloning, Expression and purification

The recombinant clones of the LILRB1 D1D2 region (residues 24–221 of the mature protein; hereafter referred to as LILRB1) and HLA-A2 were prepared as previously reported (18). High levels of pHLA-A2 complexes (comprising residues 25–300 of the mature A2 heavy chain, non-covalently associated with β_2_M and peptide) and LILRB1 were produced using conventional methods involving expression in *Escherichia coli* and *in vitro* dilution refolding (19). Renatured LILRB1 and pHLA-A2 complexes were concentrated independently, and purified by size-exclusion chromatography using a Superdex 200 column.

### 2.2 Crystallisation, Data Collection and Processing

HLA-A2 molecules in complex with non-P4 phosphorylated and non-phosphorylated epitopes were screened against commercially available crystallisation conditions with the Mosquito nanolitre robot (TTP Labtech) using the vapour diffusion method (**Table 1**). Alternative crystallisation strategies involving LILRB1 were performed using a 1:1 stoichiometric mixture of purified LILRB1 and pHLA-A2 at 10-14 mg/ml. Diffraction-grade crystals of the LILRB1-pHLA-A2 complexes appeared after 1-2 weeks at 23 °C (**Table 2**).

**Table 1.**
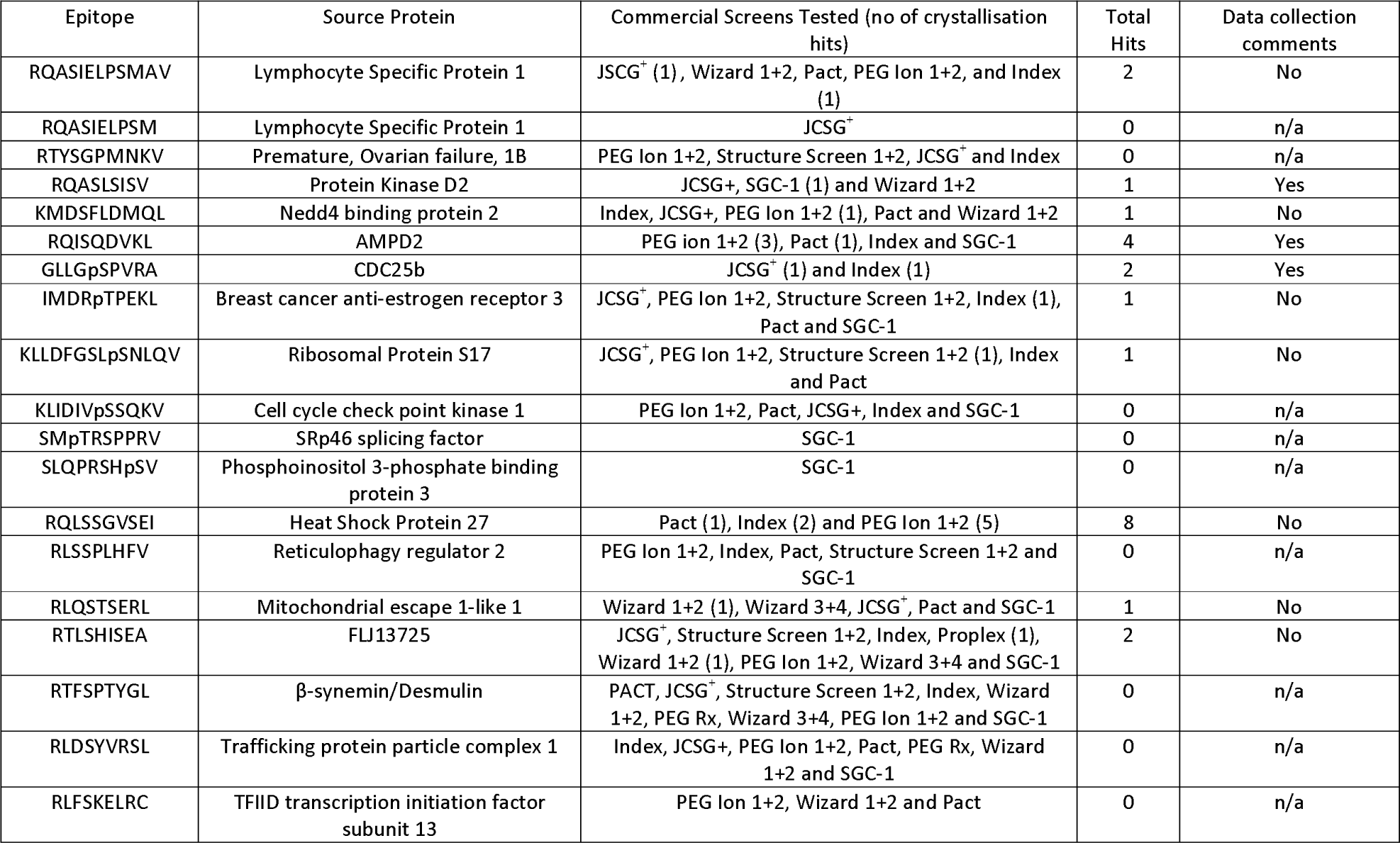
Crystallisation trials for HLA-A2 molecules bound to non-canonical or non-phosphorylated peptides.

**Table 2.** Crystallisation trials for HLA-A2 molecules bound to non-canonical or non-phosphorylated peptides in the presence of LILRB1.

| Epitope | Source Protein | Commercial Screens Tested (no of crystallisation hits) | Total Hits | Data collection comments |
| --- | --- | --- | --- | --- |
| RQASIELPSMAV | Lymphocyte Specific Protein | Proplex (8) and PAH (1) | 9 | Crystals diffracted to 2.7Å |
| RTYSGPMNKV | Premature, Ovarian failure, 1B | PAT (1) | 1 | No |
| IMDRpTPEKL | Breast cancer anti-estrogen receptor 3 | PAT/PAH (1) | 1 | Yet to be optimised |
| KLIDIVpSSQKV | Cell cycle check point kinase 1 | PAT (1) | 1 | Yet to be optimised |
| KLLDFGSLpSNLQV | Ribosomal Protein S17 | PAT | 0 | n/a |
| KMDSFLDMQL | Nedd4 binding protein 2 | PAT/PAH (2), PEG Rx (3), JCSG<sup>+</sup> (2), PEG lon 1+2 (4) and Structure Screen 1+2 (1) | 12 | Yet to be optimised |
| RLSSPLHFV | Reticulophagy regulator 2 | PAT/PAH (1) and PEG lon 1+2 (20) | 21 | Crystals diffracted to 3.2Å |
| RTFSPTYGL | β-synemin/Desmulin | PAT/PAH (1), PEG Rx (2), JCSG<sup>+</sup> (2) and PEG lon 1+2 (5) | 10 | Crystals diffracted to 2.3Å |
| RLQSTSERL | Mitochondrial escape 1-like 1 | PEG lon 1+2 | 13 | Crystals diffracted to 2.8Å |

Prior to X-ray data collection LILRB1-pHLA-A2 complex crystals were soaked in reservoir solution incorporating increasing concentrations of ethylene glycol (18-22%) and flash cooled in liquid nitrogen. X-ray diffraction data for the LILRB1-HLA-A2^ILKEPVHGV^ complex were collected to 2.4Å resolution with the ADSC Quantum 4 detector at beamline ID14-4 (ESRF). The LILRB1-HLA-A2^ILKEPVHGV^ complex crystallised in the trigonal space group P3_2_21, with two molecules per asymmetric unit, and unit cell parameters a=b=116.2Å and c=192.8Å. For all other LILRB1-pHLA2-A2 complexes, X-ray data were collected with an ‘in-house’ MicroMax 007HF rotating anode Rigaku X-ray generator using a Saturn 944 CCD detector. The LILRB1-pHLA-A2 complex typically crystallizes in the trigonal space group P3_2_21, with 2 molecules per asymmetric unit. All data were processed using the XDS suite (20) and the relevant statistics are listed in Table 3.

**Table 3.** Data processing and refinement statistics for the LILRB1-HLA-A2^ILKEPVHGV^, LILRB1-HLA-A2^RTFSPTYGL^ and LILRB1-HLA-A2^RLSSPLHYV^ complex structures.

|  | LILRB1-HLA-A2-ILK | LILRB1-HLA-A2-RTFS | LILRB1-HLA-A2-RLSS |
| --- | --- | --- | --- |
| PDB ID code | 6EWA | 6EWO | 6EWC |
| Peptide Sequence | ILKEPVHGV | RTFSPTYGL | RLSSPLHYV |
| **Data Processing** |  |  |  |
| Resolution (Å) | 48.6-2.4 (2.5-2.4) | 20-2.3 (2.4-2.3) | 20-3.2 (3.1-3.2) |
| Unit cell dimensions (Å) | 116.1, 116.1, 192.8 | 116.3, 116.3, 192.6 | 117.5, 117.5, 203.7 |
| Space Group | P3<sub>2</sub>21 | P3<sub>2</sub>21 | P3<sub>2</sub>21 |
| Total reflections | 578024 (80872) | 758810 (41698) | 149740 (12971) |
| Unique reflections | 60013 (8505) | 66969 (7259) | 26415 (2338) |
| Multiplicity | 9.6 (9.5) | 11.3 (5.5) | 5.7 (5.5) |
| Completeness (%)<sup>a</sup> | 99.6 (99.7) | 99.1 (94) | 96 (98.6) |
| R<sub>merge</sub> (%)<sup>b</sup> | 12.2 (53.7) | 10 (73.4) | 17.3 (44.2) |
| I/σ(I) | 5.2 (1.4) | 23.6 (2.8) | 10.7 (3.9) |
| **Refinement** |  |  |  |
| Resolution (Å) | 48.6-2.4 | 19.7-2.3 | 19.58-3.2 |
| Reflections used | 56939 | 63567 | 26414 |
| R<sub>cryst</sub> (%)<sup>c</sup> | 22.8 | 23.4 | 20.6 |
| R<sub>free</sub> (%)<sup>d</sup> | 27.9 | 27.6 | 24.5 |
| Protein residues | 1101 | 1117 | 1122 |
| Water molecules | 45 | 228 | - |
| **Model Geometry** |  |  |  |
| Ramachandran Plot |  |  |  |
| Most favoured | 90.8 | 89 | 88.5 |
| Additionally allowed | 8.1 | 9.6 | 10.1 |
| Generously allowed | 0.7 | 1.0 | 0.9 |
| Disallowed | 0.4 | 0.4 | 0.5 |
| RMS deviations |  |  |  |
| Bond lengths (Å) | 0.008 | 0.008 | 0.008 |
| Bond angles (°) | 1.26 | 1.29 | 1.19 |

### 2.3 Structure Determination and Refinement

The high resolution LILRB1-HLA-A2^ILKEPVHGV^ complex structure was solved by molecular replacement using MOLREP (21). The search model consisted of the LILRB1-HLA-A2^ILKEPVHGV^ complex refined to 3.4Å resolution ((18); PDB code 1P7Q). The LILRB1-HLA-A2^RQASIELPSMAV^, LILRB1-HLA-A2^RTFSPTYGL^ and LILRB1-HLA-A2^RLSSPLHFV^ complex structures were also determined by molecular replacement using the high-resolution LILRB1-HLA-A2^ILKEPVHGV^ structure complex as the search model with the co-ordinates of the ILK peptide moiety omitted. The structures were refined by alternating cycles of energy-minimization and *B*-factor refinement using CNS and REFMAC5 (22, 23). Manual rebuilding was performed with the graphics program COOT (24). All of the complexes demonstrated unequivocal F_o_-F_c_ difference density for the epitopes, which were directly built into each of the structures. The stereochemical and refinement parameters are listed in Table 3. Structure validation and analysis were carried out with CCP4 suite (25). The atomic coordinates and structure factors have been deposited in the RCSB Protein Data Bank. Figures were generated using the programs POVSCRIPT (26), Pov-Ray (http://www.povray.org) and PyMOL (27).

## 3 Results

### 3.1 HLA-A2 bound phosphopeptides can be refractory to crystallisation

Post-translationally modified peptides have emerged as an important group of antigens relevant to both autoimmunity and cancer. Phosphorylated peptides are increasingly recognised as promising tumour-associated antigens (11, 28–30) and recent studies have focused on establishing the molecular ground rules for phosphopeptide presentation by class I MHC molecules (9, 10). Our own initial molecular studies in this area, which focussed on peptides bearing phosphorylations at P4 (so called “canonical” phosphorylations, the most prevalent in the HLA-A2-restricted phosphopeptide repertoire), outlined clearly how the P4 phosphate moiety can mediate energetically significant contacts to positively charged MHC residues, while remaining highly prominent within the antigen-binding groove, and available for TCR recognition. Based on these findings the phosphate was defined as a novel “phosphate surface anchor” (9).

Subsequent to these studies, we sought to address two outstanding questions in phosphopeptide immunology: firstly, how conformationally distinct phosphopeptide antigens are compared to their non-phosphorylated counterparts (11) – an issue highly relevant for therapeutic targeting of phosphopeptide antigens, and secondly, how peptides bearing phosphorylations as positions other than P4 are accommodated in the MHC antigen binding groove – about which only very limited structural data are available. We prioritised structural studies on a range of specific pMHC complexes to address these questions, which focussed on both non-phosphorylated counterparts of previously structurally analysed P4 phosphopeptides and phosphopeptides bearing “non-canonical” (i.e. non-P4) phosphorylations.

Whereas canonical P4-phosphorylated phosphopeptides were amenable to crystallisation, the majority of their unmodified counterparts, with the exception of a few isolated examples (10, 11), proved highly intransigent to crystallisation. Similarly, structural determination of pMHC in complex with “non-canonical” (ie non-P4 phosphorylated) phosphopeptides was also hampered by the majority of such complexes being refractory to crystallisation (Table 1). Hence our attempts at structure determinations of both non-phosphorylated pMHC and non-canonical phosphopeptide antigens highlighted the need for an alternative strategy to aid pMHC crystallisation.

### 3.2 Validating the LILRB1 strategy for crystallising intransigent HLA-A2 molecules

We explored the possibility of co-crystallising intransigent pMHC complexes with a natural immune receptor ligand. One candidate receptor that reproducibly co-crystallises with HLA-A2 is LILRB1, which binds to the non-polymorphic regions of the MHC protein comprised of the α3 and β_2_M domains. Crucially, the LILRB1-pMHC interface is located distal to the peptide-binding site (18), suggesting that it is highly unlikely to interfere with epitope conformation. Comparison of the LILRB1-HLA-A2^ILKEPVHGV^ complex (18) with previous structural analyses of HLA-A2^ILKEPVHGV^ (31) failed to note any differences in the HLA-A2 bound peptide in the presence/absence of LILRB1 (18). However, LILRB1-HLA-A2^ILKEPVHGV^ structural data were only available to 3.4Å, limiting detailed analysis of the peptide conformation. To definitively resolve whether the binding of LILRB1 to HLA-A2 affected peptide conformation, we determined a higher resolution structure of the LILRB1-HLA-A2^ILKEPVHGV^ complex (to 2.4Å resolution (**Figure 1a**)), which enabled a more accurate structure of the ILK peptide moiety (**Figure 1b**). Structural overlay comparisons of this higher resolution LILRB1 /HLA-A2 structure with the HLA-A2^ILKEPVHGV^ determined in the absence of LILRB1 (31) demonstrated that the peptide binding platform in both complexes was very similar with an r.m.s.d value of 0.6Å (**Figure 1c**). Most crucially, no significant changes in structure of the ILK peptide epitope were evident upon LILRB1 binding to HLA-A2 as demonstrated by the low r.m.s.d value of 0.24Å (**Figure 1d**). This confirmed that co-crystallisation of LILRB1 with HLA-A2 complex does not alter the conformation of the MHC-bound antigenic peptide, and established a basis for exploring its potential as a chaperone for facilitating crystallisation of pMHC complex molecules.

**Figure 1.**
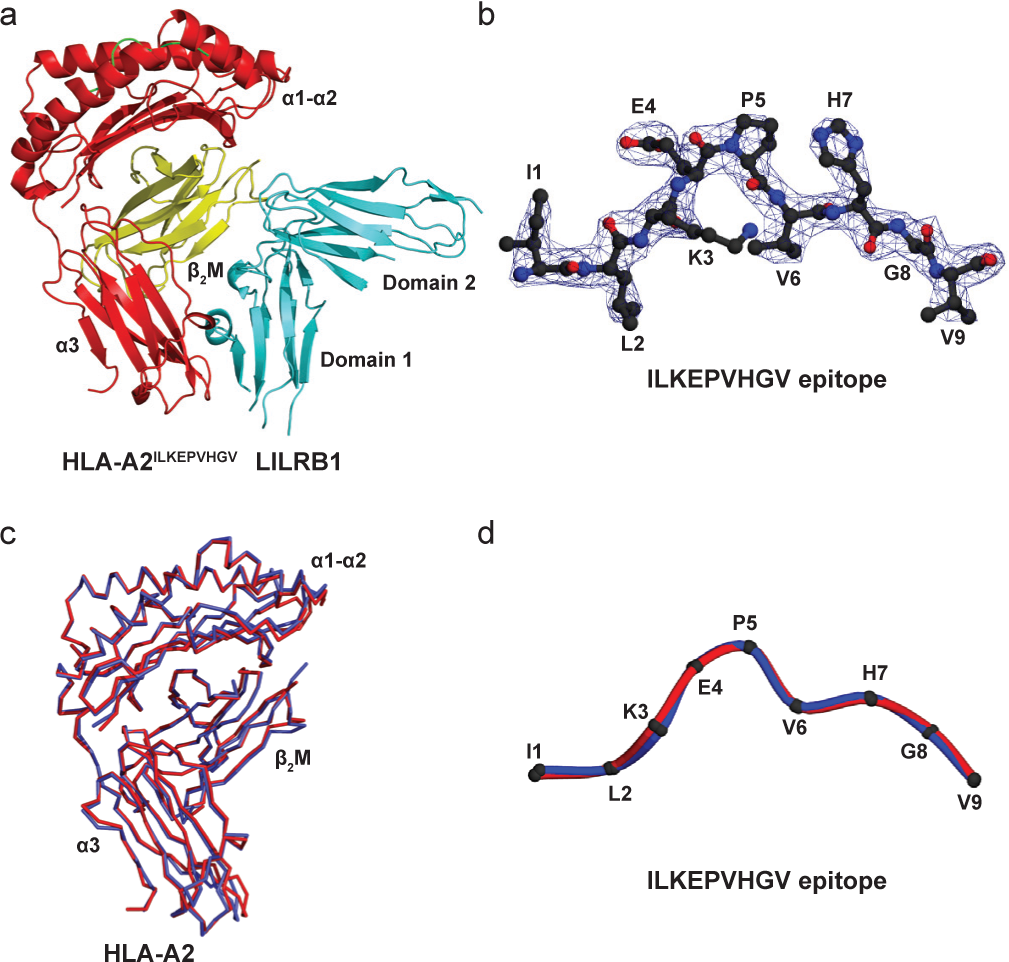
Co-crystallisation of LILRB1 with HLA-A2 does not alter the conformation of the MHC-bound antigenic peptide. (a) Ribbon representation of the LILRB1-HLA-A2^ILKEPVHGV^ Complex structure determined to 2.4Å resolution (HLA-A2 α chain (red), β2-microglobulin (yellow) and LILRB1 (cyan). (b) *2Fo-Fc* electron density map contoured at 1.0 σ (blue wire) for the ILK peptide moiety bound within the HLA-A2 peptide binding cleft. (c) Superimposition of the HLA-A2 C-α chains determined in the presence (red) and absence of LILRB1 (blue). (d) Overlay of the ILK peptide moiety derived from HLA-A2 in the presence (red) and absence (blue) of LILRB1.

### 3.3 LILRB1 facilitates crystallisation and structure determination of tumour-associated pHLA-A2-complexes

To assess whether LILRB1 could promote the crystallisation of pMHC complexes, we selected several pHLA-A2 complexes that had previously proven to be refractory to crystallisation, based on extensive nanolitre-scale crystallisation trials using commercial screening kits, at concentrations commonly used for class I MHC crystallisation (typically 10-25 mg/ml). These were generally tumour associated phosphopeptides, and their unphosphorylated counterparts (32). LILRB1 and pHLA-A2 complexes were produced as previously described (18).

Initial attempts at crystallising pMHC complexes previously found to be refractory to crystallisation alone, frequently resulted in multiple hits in co-crystallisation trials with LILRB1 (**Table 2**). Initial attempts focussed on non-phosphorylated antigens, which involved equilibrating against conditions that had yielded LILRB1-HLA-A2^ILKEPVHGV^ crystals, revealed a crystallisation solution (PEG 3350, Ammonium acetate and, Tris-HCl – hereafter referred to as PAT) that proved somewhat generic, as it was successful in providing useful primary hits for a diverse subset of pMHC complexes previously intransigent to crystallisation. An example included the 10-mer HLA-A2^KMDSFLDMQL^ peptide complex, crystals of which grew with a morphology similar to that of LILRB1-HLA2^ILKEPVHGV^ crystals (**Figure 2a**), in the presence of LILRB1. In addition, the 12-mer HLA-A2^RQASIELPSMAV^ complex, also previously intransigent to crystallisation, yielded crystals with LILRB1 that grew in an optimised form of the generic PAT crystallisation reagent (comprised of 20% PEG 3350, 0.2M ammonium acetate and 0.1M HEPES pH 7.4 – hereafter referred to as PAH) (**Figure 2b**). Crucially, this same PAH condition failed to crystallise HLA-A2^RQASIELPSMAV^ in the absence of LILRB1, underlining the critical chaperone function of LILRB1 in the crystallisation process. Importantly, the PAT condition also demonstrated considerable promise for crystallising HLA-A2 molecules bound to non-canonical phosphopeptides, including the 9-mer (IMDRpTPEKL) (**Figure 2c**) and 11-mer (KLIDIVpSSQKV) (**Figure 2d**).

**Figure 2.**
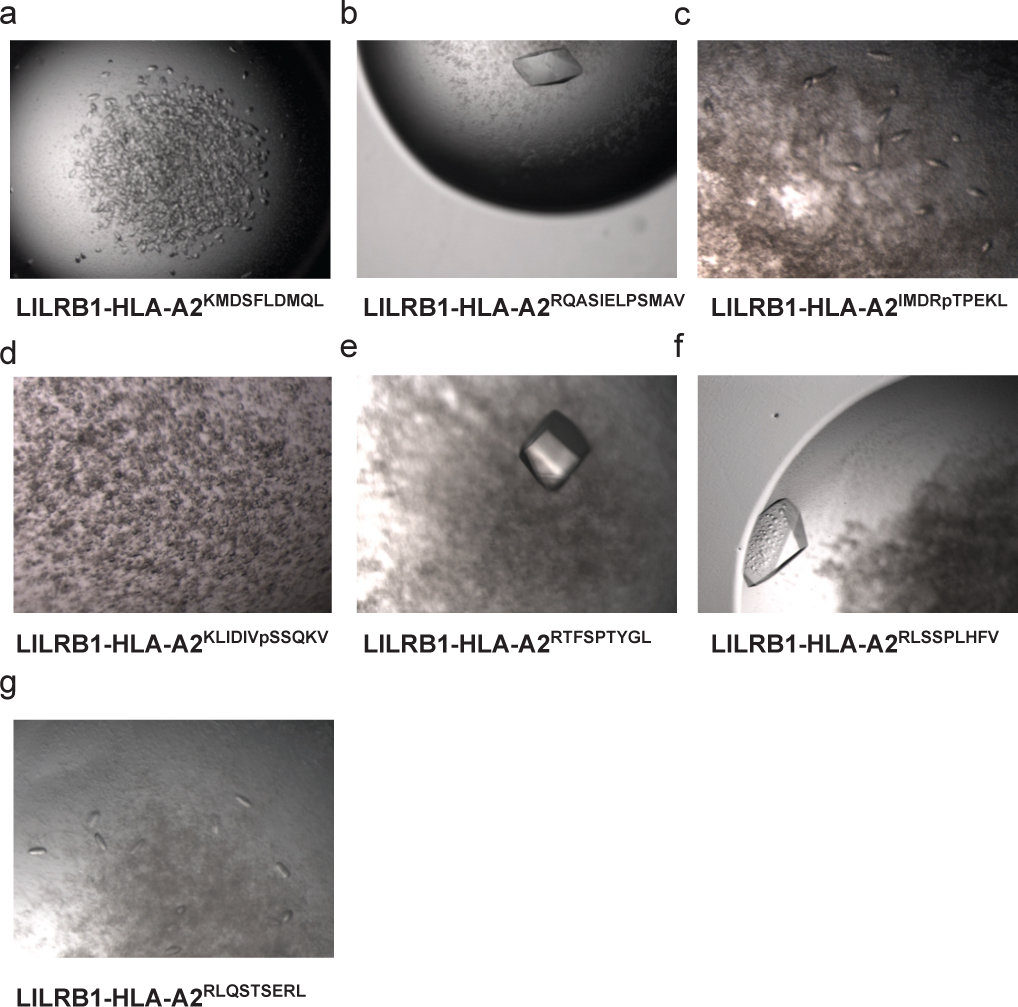
Crystallisation of intransigent HLA-A2-peptide complexes with LILRB1. Crystal morphologies of LILRB1-HLA-A2^KMDSFLDMQL^ (a), LILRB1-HLA-A2^RQASIELPSMAV^ (b), LILRB1-HLA-A2^IMDRpTPEKL^ (c), LILRB1-HLA-A2^KLIDIVpSSQKV^ (d), LILRB1-HLA-A2^RTFSPTYGL^ (e), LILRB1-HLA-A2^RLSSPLHFV^ (f) and LILRB1-HLA-A2^RLQSTSERL^ (g).

Despite successful use of the PAT condition as a generic crystallisation condition for a subset of pMHC, for other pMHC complexes fresh crystallisation hits were identified in the presence of LILRB1 following rescreening of complexes against commercial sparse matrix kits. An example was HLA-A2^RTFSPTYGL^ crystals, which yielded co-crystals with LILRB1 from the PEG Ion screen in the presence of 0.2M potassium sodium tartrate and 20% PEG 3350 (**Figure 2e**). A similar LILRB1 co-crystallisation screening strategy for the HLA-A2^RLSSPLHFV^ complex resulted in initial micro-crystals obtained in drops equilibrated against 3% Tacsimate pH 6 and 12.8% PEG 3350, after which further optimisations in the presence of dimethyl sulphoxide produced large well ordered LILRB1-HLA-A2^RLSSPLHFV^ complex crystals (Figure 2f). Finally, it was possible to crystallise the HLA-A2^RLQSTSERL^ complex, which we found previously was intransigent to crystallisation attempts, in complex with LILRB1 in the presence of 0.2M Potassium Acetate and 20% PEG 3350, resulting in microcrystals worthy of further optimisation (**Figure 2g**).

When combining the two groups of antigens we focussed on (non-phosphorylated counterparts of P4 phosphopeptides, and non-canonical phosphopeptides), only 10 out of the 19 pMHC complexes yielded hits with conventional trials (Table 2). In contrast, a majority of pMHC complexes (8/9) generated hits when co-crystallised with LILRB1 (Table 3). Furthermore, several complexes yielded multiple independent hits thereby increasing the likelihood of growing diffraction-grade crystals (>5, Table 3).

Crystals produced using the LILRB1 co-crystallisation strategy were of sufficient quality for data collection. Optimised LILRB1 co-crystals of the unmodified 12-mer, LILRB1-HLA-A2^RQASIELPSMAV^ complex (**Figure 2d**), permitted data collection and structure determination to 2.7Å (11). Moreover, LILRB1 co-crystals of the HLA-A2^RTFSPTYGL^ and HLA-A2^RLSSPLHFV^ complexes diffracted X-rays to 2.4Å and 3.2Å, resulting in full structure determinations (**Figure 3**). The quality of the resulting electron density maps were significantly improved using two-fold non-crystallographic symmetry averaging, which is present in all LILRB1-pHLA-A2 complex crystals, thereby aiding model building and structure determination (**Figure 3**). Collectively, these results clearly highlight the potential of exploiting LILRB1 as a crystallisation chaperone, to facilitate X-ray crystallographic analyses of biologically important pMHC complexes.

**Figure 3.**
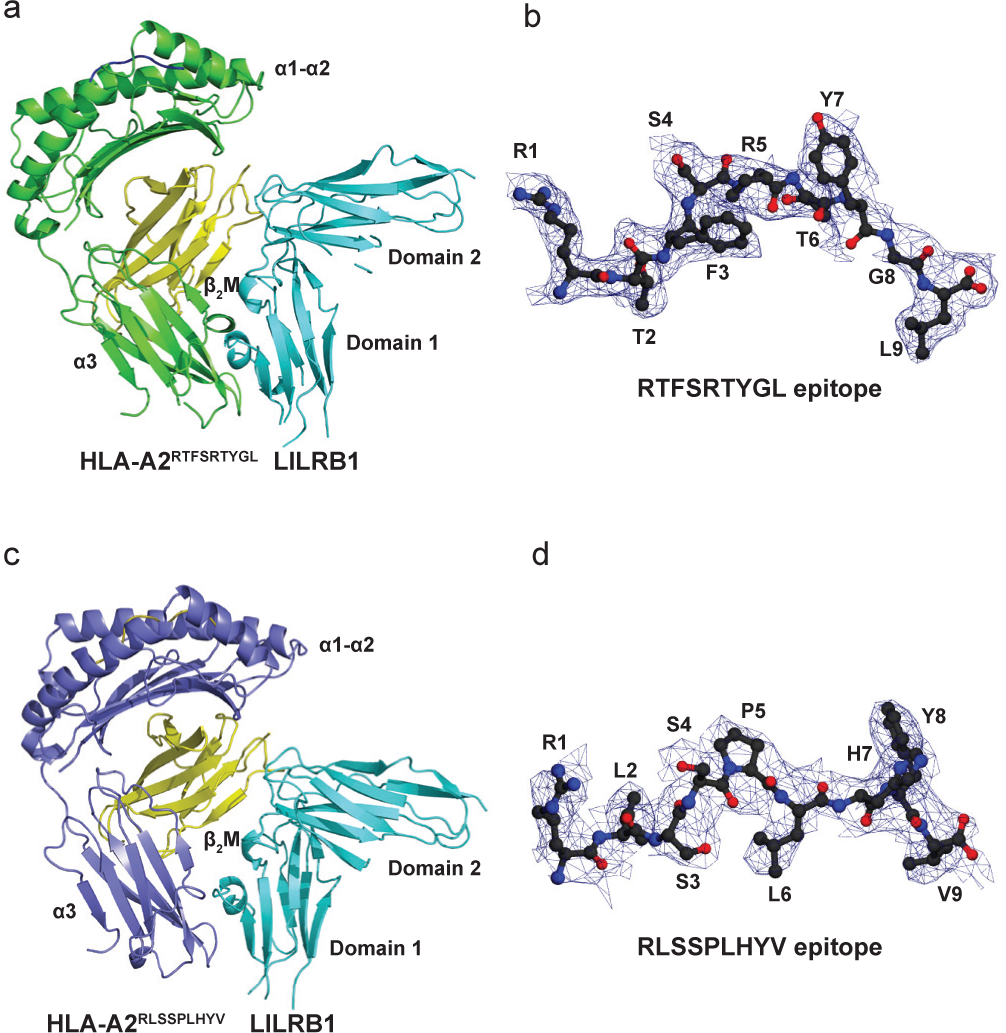
Crystal structures of LILRB 1-HLA-A2 ^RTFSPTYGL^ and LILRB1-HLA-A2^RLSSPLHFV^ complexes. (a) Ribbon representation of the LILRB1-HLA-A2^RTFSPTYGL^ complex structure determined to 2.3Å resolution (HLA-A2 α chain (green), β2-microglobulin (yellow) and LILRB1 (cyan). (b) *2Fo-Fc* electron density map contoured at 1.0 σ (blue wire) for the RTF peptide moiety bound within the HLA-A2 peptide binding groove. (c) Ribbon representation pf the LILRB1-HLA-A2^RLSSPLHFV^ complex structure determined to 3.2Å resolution (HLA-A2 α chain (purple), β2-microglobulin (yellow) and LILRB1 (cyan). (d) *2Fo-Fc* electron density map contoured at 1.0 σ (blue wire) for the RLS peptide moiety bound within the HLA-A2 peptide binding cleft.

## 4. Discussion

Structural studies of class I peptide MHC structures continue to make major contributions to our understanding of important areas of immunobiology. However, despite availability of numerous pMHC structures, reliable structural analyses of predefined pMHC targets is still challenging, as certain pMHC complexes can be intractable to crystallisation. This represents a significant impediment to molecular studies aiming to define the role of MHC-restricted antigenic peptide epitopes in specific immunobiological contexts such as disease pathogenesis and immunotherapeutic development. In the context of MHC alleles that have been crystallised, this phenomenon is superficially surprising, given conservation of the alpha chain α3 and β_2_M, and the fact that only the peptide moiety would be altered between each individual pMHC complex. Whilst the molecular basis underlying it is unclear, it is likely to result from the hugely diverse properties of bound peptides. Given the strong link between protein stability and propensity for crystallisation, one significant factor is likely to be the wide span of peptide binding affinities for MHC, and the relative kinetics of complex dissociation and aggregation, versus crystal nucleation. However, our demonstration that peptides with similar epitope sequence and binding affinities, such as RQA_V in its phosphorylated and non-phosphorylated states (11), may not exhibit the same propensity for crystallisation, suggests that factors other than peptide affinity, such as the potential of peptide conformation to favour or disrupt crystal packing interactions, or differential complex solubility, are likely to be relevant to crystal formation.

In this study we investigated a novel strategy for circumnavigating crystallisation of intransigent pMHC complexes. The approach relies upon the addition of a natural ligand of class I MHC, LILRB1, to promote alternative, and in many cases more optimal crystal packing contacts. Our findings, focused in this study on the HLA-A2 allele, highlight that LILRB1 can serve as an effective non-covalent crystallization chaperone for pMHC complexes. This strategy offers several advantages. Firstly, since co-crystallisation with LILRB1 does not perturb the biologically critical α1α2 peptide-binding platform, it allows *bone fide* peptide conformation to be observed. Secondly, the approach is experimentally highly feasible. LILRB1 is easily over-expressed in large amounts into *E. coli* inclusion bodies (typical yields of 100g/l), and renaturation and purification is relatively efficient. Moreover, crystallisation optimisation with LILRB1, which exploits a generic crystallisation condition in many cases, is extremely efficient, and results in the production of large crystals within a relatively short time interval (<2 weeks), often of a sufficient size for data collection. Moreover, while such crystals yield acceptable data using ‘in-house’ sources, clearly use of synchrotron sources would inevitably improve resolution further. In addition, the availability of the higher resolution structure of LILRB1 provides useful model-based phase information necessary for resolving LILRB1-pHLA-A2 complexes, a process that has become increasingly routine since all LILRB1-pHLA-A2 crystals exhibit similar unit cell constants, even if grown in chemically distinct conditions. Typically, the presence of two LILRB1-HLA-A2-petide complexes in the asymmetric unit allows non-crystallographic symmetry averaging, improving the quality of the electron density. Thirdly, based on the evidence we present here, LILRB1 co-crystallisation is clearly an approach capable of catalyzing crystallisation of a diverse range of peptides in the context of HLA-A2, including those previously intransigent to crystallisation. Finally, in view of the fact that LILRB1 recognises the non-polymorphic regions of classical and non-classical pMHC molecules, there is considerable potential for extending this strategy to facilitate crystallisation of a diverse range of class I MHC molecules.

Development of LILRB1 as a crystallisation chaperone for pMHC could have several applications. Immune presentation and recognition of post-translationally modified peptide antigens is increasingly recognised as an area of immunobiological importance, not least in the context of cancer immunosurveillance and immunotherapy. We have successfully applied the method to dissect the effects of phosphorylation of peptide conformation. Of relevance in this context, so-called “non-canonical” phosphopeptide HLA-A2 complexes, for which limited structural data are available, have proven to be relatively intransigent to conventional crystallisation attempts; furthermore unmodified counterparts of naturally occurring phosphopeptides tend to be notably lower affinity, and would be expected to represent challenging crystallisation targets. Use of LILRB1 as a crystallisation chaperone facilitated crystallisation of several such peptides. In addition, the method may also be particularly suitable for longer, more bulged peptides (either unmodified or those bearing bulky post-translational modifications), where conventional class I MHC crystal packing interactions may be disrupted. Of relevance to this grouping, exhaustive conventional attempts to crystallise the bulky 12-mer unmodified HLA-A2^RQASIELPSMAV^ complex failed entirely, despite in this case an equivalent affinity to the naturally phosphorylated form. The LILRB1 chaperone approach quickly led to its structure determination, allowing us to demonstrate that phosphorylation of this leukaemia-associated epitope resulted in an unprecedented conformational change relative to this unmodified form, creating a highly distinct conformational “neoepitope” (11). Indeed, examination of the structure of unphosphorylated HLA-A2^RQASIELPSMAV^ in complex with LILRB1 provided a molecular explanation for its failure to crystallise alone, highlighting a more pronounced bulge to the peptide conformation at P8 (Proline) that precluded crystallisation in the same mode as the phosphorylated form (HLA-A2^RQApSIELPSMAV^) by causing steric clashes with a neighbouring molecule. This observation highlights that altered crystal contacts introduced by LILRB1 co-crystallisation can clearly circumvent such problems. A second scenario, peptide anchor modification, which is an immunotherapeutic approach used to boost antigen immunogenicity whereby peptide immunogens are engineered with modified anchor residues to optimise MHC binding, is another setting where the LILRB1 crystallisation chaperone methodology could be productively applied. Here the intention is to increase MHC affinity but without altering peptide conformation presented to TCR. Structural comparisons of unmodified and modified forms (the former by definition of low affinity) are likely to be highly informative in this setting.

In light of our results, we propose that other class I MHC receptors could be exploited as alternative crystallisation chaperones – for class I pMHC, the two most likely candidates are LILRB2 and CD8αα, both of which bind to a broad range of class I MHC molecules (17, 33). Moreover, previous structural studies of both LILRB2 and CD8αα immune receptors in complex with class I MHC have highlighted that they interact with sites of the MHC that are distal to the antigen binding platform and therefore are highly unlikely to influence epitope conformation (34, 35). LILRB2 displays an overlapping but distinct MHC-I recognition mode relative to LILRB1 and predominantly mediates hydrophobic contacts to the HLA-G α3 domain (35). Moreover, structural comparisons of HLA-G and its bound peptide in the presence and absence (36) of LILRB2 have demonstrated no substantial shifts in conformation (**Figures 4 a-b**) thus confirming the potential of LILRB2 as a tool for promoting protein crystallisation of non-classical MHC molecules. In contrast, the CD8αα-MHC binding interaction mode significantly differs to that of LILRB1 and LILRB2 forming interactions with the α2 and α3 domains of HLA-A2 as well as β_2_M (34), but similarly has no significant effects on the conformation of the α1 α2 peptide binding platform (**Figures 4 c-d**). Therefore LIR-2 and CD8αα could have potential as crystallisation chaperones for pMHC, although it is unclear whether these will provide advantages to the current strategy. In summary, the success we have observed with the LILRB1 co-crystallisation approach suggests that this method offers an effective means for promoting crystallisation of intransigent pMHC complexes. We predict that co-crystallization of pMHC molecules with LILRB1 will be a valuable addition to the growing repertoire of tools available to resolve the macromolecular crystallisation bottleneck for class I pMHC molecules.

**Figure 4.**
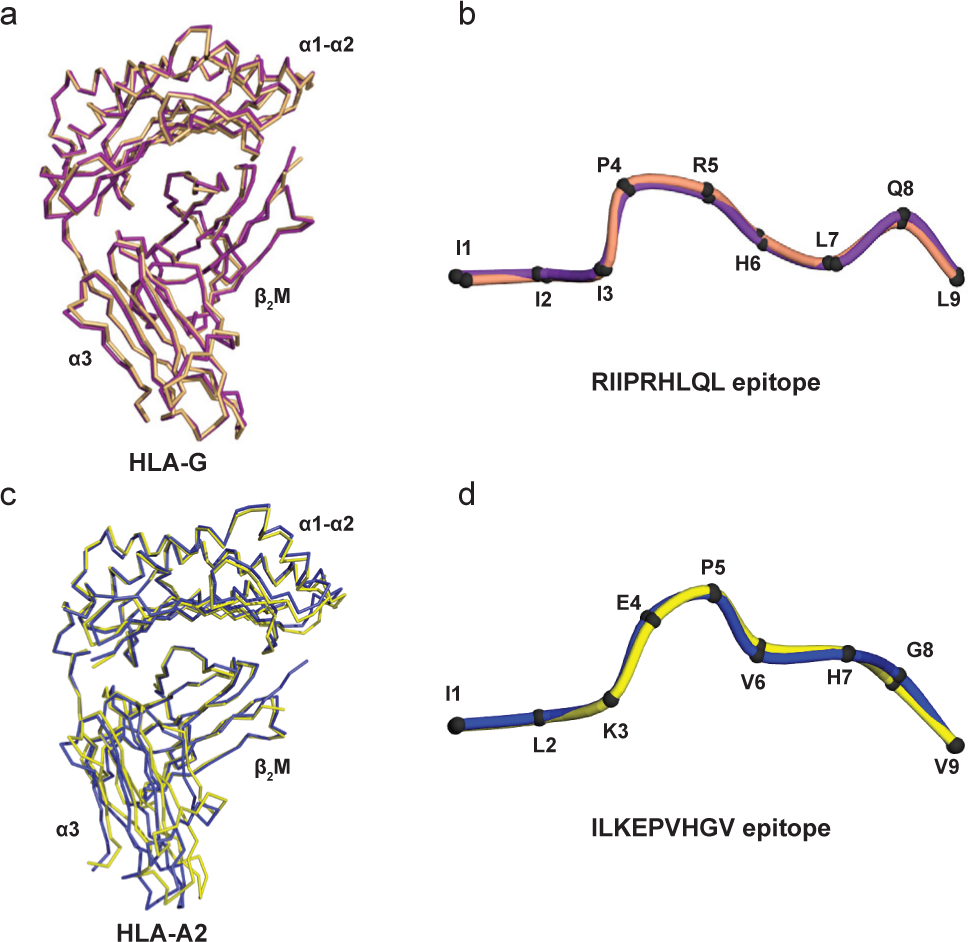
Co-crystallisation of LILRB2 or CD8αα with MHC class I molecules does not affect the conformation of the bound antigenic peptide. (a) Superimposition of the HLA-G C-α chains determined in the presence (purple) and absence (pink) of LILRB2. (b) Overlay of the RII peptide moiety derived from HLA-G in the presence (purple) and absence (pink) of LILRB2. (c) Superimposition of the HLA-A2 C-α chains determined in the presence (yellow) and absence (blue) of CD8αα. (d) Overlay of the ILK peptide moiety derived from HLA-A2 in the presence (yellow) and absence (blue) of CD8αα.

## Author contributions

FM and DHS designed the study and carried out experiments. FM and DHS analysed data and wrote the manuscript. BEW designed the study, analysed data and wrote the manuscript.

## Acknowledgements

We acknowledge the Birmingham Protein Expression Facility for assistance with recombinant protein production. This work was funded by a Wellcome Trust New Investigator Award. DHS was supported by a Medical Research Council studentship.

## Conflict of interest

The authors declare that they have no conflicts of interest with the contents of this article.

